# Paired associated SARS-CoV-2 spike variable positions: a network analysis approach to emerging variants

**DOI:** 10.1101/2023.05.04.539462

**Authors:** Yiannis Manoussopoulos, Cleo Anastassopoulou, John P. A. Ioannidis, Athanasios Tsakris

## Abstract

Amino acids in variable positions of proteins may be correlated, with potential structural and functional implications. Here, we apply exact tests of independence in R × C contingency tables to examine noise-free associations between variable positions of the SARS-CoV-2 spike protein, using as a paradigm sequences from Greece deposited in GISAID (N=6,683/1,078 full-length) for the period February 29, 2020 to April 26, 2021 that essentially covers the first three pandemic waves. We examine the fate and complexity of these associations by network analysis, using associated positions (exact p≤0.001 and Average Product Correction ≥2) as links and the corresponding positions as nodes . We found a temporal linear increase of positional differences and a gradual expansion of the number of position associations over time, represented by a temporally evolving intricate web, resulting in a non-random complex network of 69 nodes and 252 links. Overconnected nodes corresponded to the most adapted variant positions in the population, suggesting a direct relation between network degree and position functional importance. Modular analysis revealed 25 *k*-cliques comprising three to 11 nodes. At different *k*- clique resolutions, one to four communities were formed, capturing epistatic associations of circulating variants (Alpha, Beta, B.1.1.318), but also Delta, which dominated the evolutionary landscape later in the pandemic. Cliques of aminoacidic positional associations tended to occur in single sequences, enabling the recognition of epistatic positions in real-world virus populations. Our findings provide a novel way of understanding epistatic relationships in viral proteins with potential applications in the design of virus control procedures.

**Importance:** Paired positional associations of adapted amino acids in virus proteins may provide new insights for understanding virus evolution and variant formation. We investigated potential intramolecular relationships between variable SARS-CoV-2 spike positions by exact tests of independence in R × C contingency tables, having applied Average Product Correction (APC) to eliminate background noise. Associated positions (exact p≤0.001 and APC≥2) formed a non-random, epistatic network of 25 cliques and 1-4 communities at different clique resolutions, revealing evolutionary ties between variable positions of circulating variants, and a predictive potential of previously unknown network positions. Cliques of different sizes represented theoretical combinations of changing residues in sequence space, allowing the identification of significant aminoacidic combinations in single sequences of real-world populations. Our analytic approach that links network structural aspects to mutational aminoacidic combinations in the spike sequence population offers a novel way to understand virus epidemiology and evolution.

## Introduction

SARS-CoV-2 spike, a class I fusion glycoprotein typically of 1,273 amino acids, mediates virus entry into cells via a multifunctional mechanism (1). Preservation of the cell entry process is ensured through amino acid conservation in spike that secures the functionality of its tertiary structure. However, as the virus needs to adapt to new microenvironments, functionality is preserved through positional alterations across the protein, and possibly beyond, at allosteric sites through epistasis (2, 3). Several changes in spike that characterize variants have been and continue to be identified as SARS-CoV-2 continues its adaptation in humans.

New variants emerge as the virus mutates in response to prevailing selection pressures through adaptations across spike (and other) proteins (4, 5). The occurrence of multilocational aminoacidic changes raises the question of potential conditional associations among altered sites and their role in virus biology. Changes may contribute jointly to spike conformations needed for simultaneous immunogenic escape and functionality preservation (6). Interactions that could have a profound effect on the biological characteristics of the virus, may concern associations of several positions at a time (7).

Here, we examine the altered positions of all spike sequences deposited in GISAID from Greece during the first three pandemic waves and develop a method based on pairwise exact tests of independence to investigate the associations between all combinations of variable positions (8–10) in relation to variant formation. We further study the complexity of aminoacidic adaptations over time by partitioning the dataset in month intervals and creating the respective evolving networks, using noise-free (MIp Z-score ≥ 2) significant associations (exact p≤0.001) as links and the corresponding positions as nodes.

We found a gradual increase of altered positions when compared to the original sequence and identified 652 unique sequences distributed in different frequencies. There was a gradual increase in the complexity of paired associations over time, depicted by a temporally developing network that resulted in a non-random intricate web of interconnected spike positions at the end of the third pandemic wave. Modular network analysis revealed tightly associated positions forming cliques and communities that hold potentially important information. Cliques represented single sequence positional associations, which could be identified in real-world sequence populations, while communities captured homogeneous positional information with potential prediction ability of unknown positions’ functionality. Mapping through network analysis the evolution of the associated variable positions in existing sequences may enable the classification of functional constellations of mutations in emerging variants.

## RESULTS

### Circulating variants

Overall, 1,090 whole genome SARS-CoV-2 sequences from Greece were retrieved from GISAID, which comprised all available records for the period February 29, 2020, to April 26, 2021. Twelve sequences were incomplete and excluded from analysis. By May 31, 2021, an additional 5,605 partial (spike) records had become available and were also obtained for analysis. Twenty four of the 5,605 records were included in the analysis with approximated date annotations. Thus, 6,683 spike protein sequences (1,078 translated from full-length nucleic acid+5,605 partial) were analyzed. According to the strict definitions (SI Table 1), the variant distribution was as follows: 69.8% (n=4,668) B.1.1.7 (Alpha), 9.01% (n=602) B.1.1.318, 0.554% (n=37) B.1.351 (Beta), 0.015% (n=1) P.1 (Gamma), 0.015% (n=1) B.1.525 (Eta), 0.015% (n=1) B.1.617.1 (Kappa), 0.299% (n=20) index virus and 20.1% (n=1,345) other, unclassified sequences (SI File 1). Although Delta was not detected based on the strict profile, eight (0.12%), presumably Delta-precursors, were detected. The dates on which variants were first detected in the country are shown in SI Table 2.

### Spike variability beyond known variant signatures

Compared to the index virus, 652 unique aminoacidic signatures were detected, defined by different combinations of changes across the spike protein at varying frequencies (SI File 1, F/H). The mean (±sem) and the median (IQR) of changes in the 6,683 sequences were 8.47 (±0.041) and 10 (4), respectively, and they did not differ by type [8.21 (±0.11) and 10 (4) in the full (n=1,078) and 8.52 (±0.04) and 10 (4) partial genome dataset (n=5,606)]. The spatiotemporal distribution of samples is shown in SI Fig.1, 2. Most sequences were obtained from the Athens metropolitan region (Attica), principally during the third wave. Only partial sequences were obtained during the second wave.

Using local regression (“loess”) and linear regression, we found a trend of increasing changes with time, from one to about ten during the first two waves (Feb 26-May 3, 2020, and Sept 1-Dec 31, 2020), to ten and up to 15 amino acid substitutions during the third wave (Feb 1- Jun 30, 2021) (Fig.1). The trend was steeper during the first two waves compared to the third. The rate 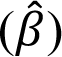 of accruing aminoacidic changes per day, between a sequence at a given time and the wild type sequence (Hamming distance), was 0.015 and 0.016 during the first and second waves, respectively, and 0.003, ∼5-fold lower, during the third wave. All rates were significant (p<0.005). Alpha was first detected towards the end of the second wave (on Dec 11, 2020) and became dominant during the third wave. During the early pandemic stages, spike sequences could differ from the index virus at a single aminoacidic position, commonly D614G (B.1) (11), a substitution that proved crucial for SARS-CoV-2 competitive fitness in terms of cell infectivity and viral transmission (12, 13). Additional aminoacidic changes accumulated over time in the ensuing sequences, often in a stepwise fashion as shown in Table 1.

**Figure 1.**
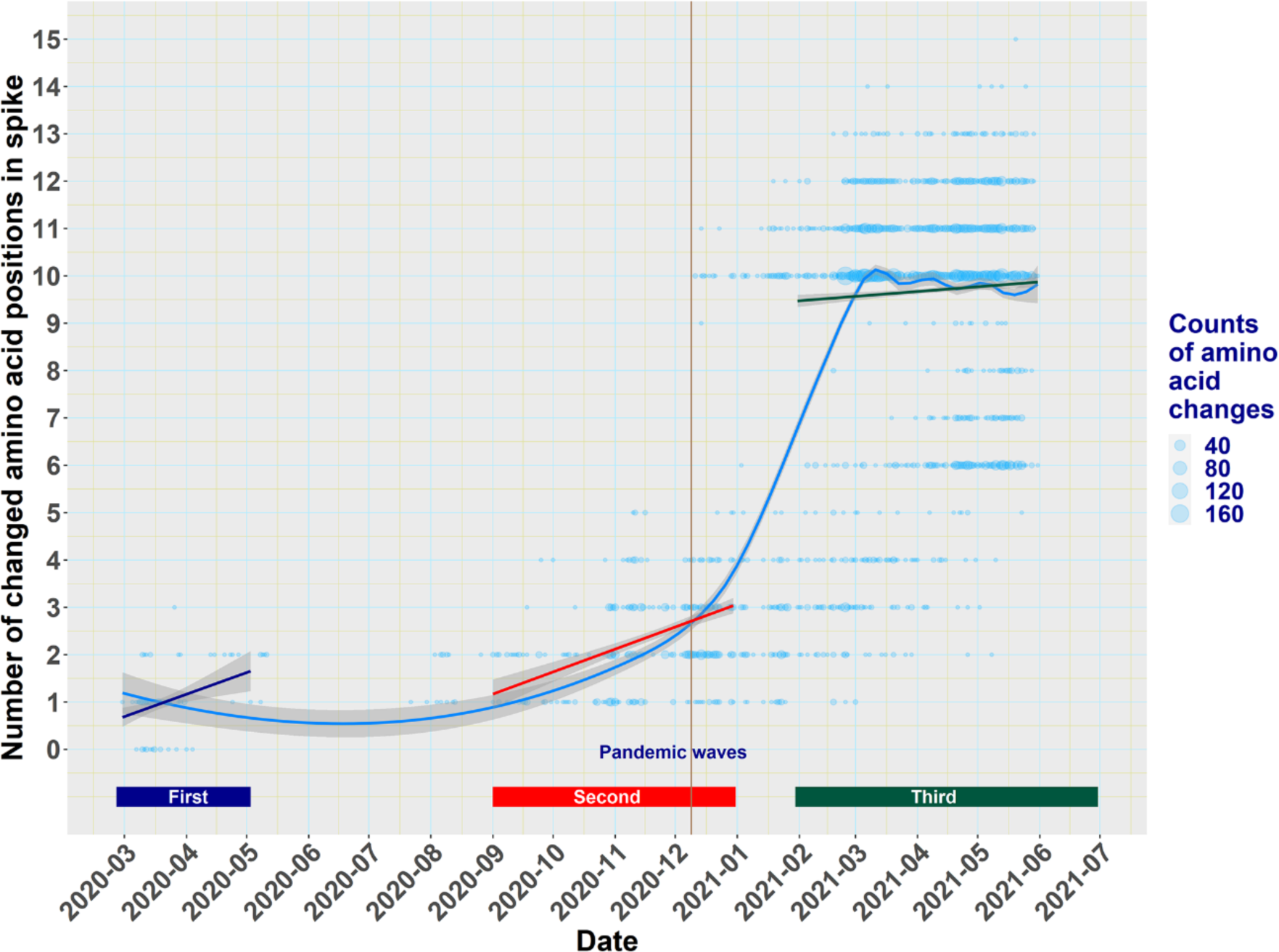
Local polynomial regression (loess) and linear regression trend lines of the number of spike changes of all unique SARS-CoV-2 isolates in relation to time. The cyan “loess” line shows the overall trend of aminoacidic changes during the three pandemic waves. The blue, red and brown regression lines show the corresponding trends during the three pandemic waves. Shaded areas across lines represent 95% confidence intervals. Dots represent unique spike sequences and dot size is analogous to the frequency of the number of changes. The vertical red line indicates the time of detection of the first Alpha variant isolate in Greece (December 11, 2020, EPI_ISL_2301679). COVID-19 pandemic waves were as follows: First wave: Feb 26 - May 3, 2020; second wave: Sept 1 - Dec 31, 2020; third wave: Feb 1 - Jun 30, 2021.

**Table 1.**
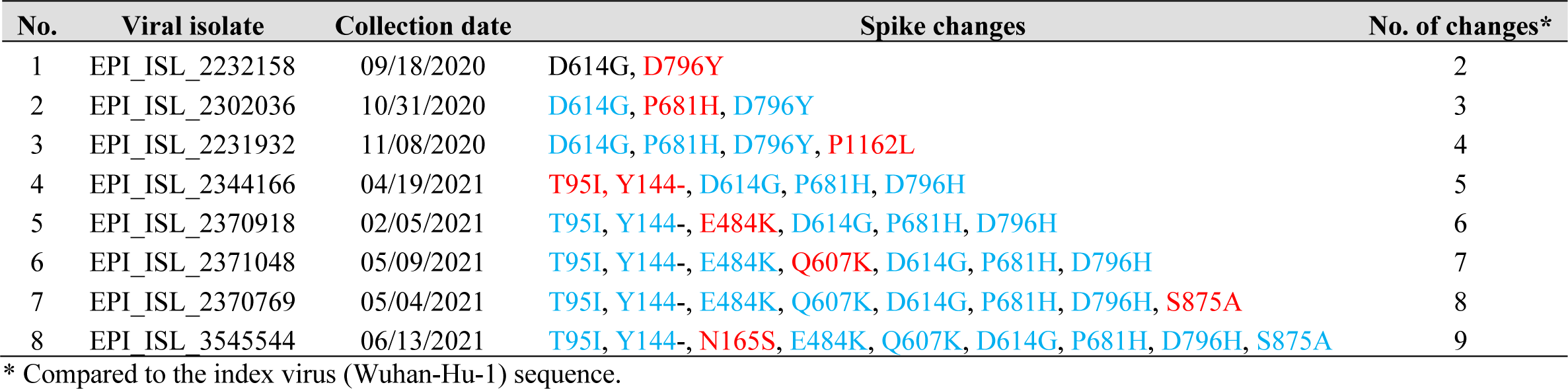
Trend of temporally sequential amino acid changes in the spike protein of SARS-CoV-2 isolates. In the early stages of the pandemic, most isolates differed from the previous sequence by a single positional adaptation. Newly added changes that often occurred in a stepwise fashion are shown in red and preserved substitutions from the previous time point are shown in blue. This stepwise progress coincided with the sequential addition of all mutated sites that formed variant Β.1.1.318, as also observed in the largest *k-*7 community of the spike network (Fig.5). Dates indicate the first detection event in Greece, as reported in the corresponding GISAID record.

| No. | Viral isolate | Collection date | Spike changes | No. of changes* |
| --- | --- | --- | --- | --- |
| 1 | EPI_ISL_2232158 | 09/18/2020 | D614G, **D796Y** | 2 |
| 2 | EPI_ISL_2302036 | 10/31/2020 | **D614G**, **P681H**, **D796Y** | 3 |
| 3 | EPI_ISL_2231932 | 11/08/2020 | **D614G**, **P681H**, **D796Y**, **P1162L** | 4 |
| 4 | EPI_ISL_2344166 | 04/19/2021 | **T95I**, **Y144-**, **D614G**, **P681H**, **D796H** | 5 |
| 5 | EPI_ISL_2370918 | 02/05/2021 | **T95I**, **Y144-**, **E484K**, **D614G**, **P681H**, **D796H** | 6 |
| 6 | EPI_ISL_2371048 | 05/09/2021 | **T95I**, **Y144-**, **E484K**, **Q607K**, **D614G**, **P681H**, **D796H** | 7 |
| 7 | EPI_ISL_2370769 | 05/04/2021 | **T95I**, **Y144-**, **E484K**, **Q607K**, **D614G**, **P681H**, **D796H**, **S875A** | 8 |
| 8 | EPI_ISL_3545544 | 06/13/2021 | **T95I**, **Y144-**, **N165S**, **E484K**, **Q607K**, **D614G**, **P681H**, **D796H**, **S875A** | 9 |

\* Compared to the index virus (Wuhan-Hu-1) sequence.

Some positional combinations were favored and appeared in higher frequencies. The most frequent simple combinations were the A222V/D614G dyad (Pango B.1.177), and the L18F/A222V/D614G triad (Pango B.1.177.44), with 71 and 223 counts, respectively (SI File 1, F/H). Overall, there were 57 unique sequences with 2-6 mutations, comprising ∼9% of the unique sequence population. Other equally or more complex patterns within the genetic context of lineages, were also evident. Dyad D1084E/G1219V was present in 22 unique sequences, scoring 190 counts in addition to the ten (most prevalent) Alpha-defining substitutions, comprising ∼3% of unique sequences. Similarly, Q607K/S875A present in two unique sequences, scored 21 counts (3%) along with the six (second most prevalent) B.1.1.318-defining changes (SI File 1, Table 1).

### Pairwise associations of variable positions

To examine the potential associations between variable positions, we analyzed further the spike protein alignment of the 1,078 translated full-length sequences available in GISAID for the study period. This subset had similar statistics to the partial genome spike sample (n= 5,605), and except for the last part of the third wave, it also had analogous spatiotemporal distribution (SI Figs. 1,2). Compared to the index virus, 1,103 conserved sites and 170 variable positions were found, the associations of which were examined by the Pearson Chi Squared exact test of independence in R × C contingency tables for all pairwise combinations.

An exemplary table examining two functional positions (numbers 681 and 501) is shown in SI Table 3. Position 681 at the S1/S2 cleavage site is likely related to fusogenicity (14), while position 501 in the receptor binding domain (RBD) is a site related to ACE2 binding (15–18). The total number of contingency tables covering all possible pairwise amino acid combinations was 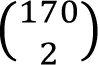 = 14,365. Most pairs 13,760 (≈ 95.8%) had p≥0.05 suggesting no association, 286 pairs (≈2.0%) had 0.001<p<0.05 suggesting moderate associations and 318 pairs (≈2.2%) had p≤0.001 suggesting strong associations between residues. In this last category, 66 pairs had 0.001≤p<0.0001 and 152 pairs had p≤10^-5^. Correction of family-wise error by the Benjamini-Hochberg method (19) indicated as significant associations those with p≤0.001 (values were rounded to four digits).

### Phylogenetic signal, random noise and recombination interference

To eliminate random background noise and phylogenetic history from shared ancestry, we calculated MIp and *Z*-scores of all pairwise combinations of 1,276 positions (1,273 + 3 insertions) in the spike alignment, yielding a total of 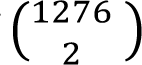 =813,450 values . Of them 314 (0.038%) pairs had *Z*-scores ≥ 2. There were 69 positions in paired associations having Z-scores ≥ 2 and all, except one (i.e., position 158), were identical to the 70 positions found by exact tests (p ≤ 0.001). Of the 318 pairs with significant associations (p ≤ 0.001), 252 (79%) had *Z*-scores ≥ 2, suggesting a high approximation of results of exact tests and MIp scores. Associated pairs of the excluded position 158 were comparable by the two methods, although the exact probability was ≤ 0.001 and *Z*- scores were < 2, ranging from 0.5 to 1.6.

To examine potential recombination events in variable positions, we applied ten recombination detection algorithms, via the RDP5 program in the nucleic acid sequences corresponding to the spike protein sequences used in the exact test. We found no recombination events among studied virus isolates.

### The intramolecular network of the spike protein

#### Network construction and basic properties

To gain insight into the complexity of relationships among variable positions in the spike protein alignment, we constructed and examined the corresponding network (20, 21) using all significant exact pair associations (p≤0.001) with Z-score ≥ 2 as links and the involved positions as nodes. The resulting network had 69 nodes and 252 links in a hub-like hierarchy in which a few nodes dominated most connections (Fig.2). The network was fragmented, holding nine independent dyads (18 nodes/nine links), two triads (six nodes/six links) and one large component (the main network) of 45 nodes and 237 links. Five of the dyad positions were found outside the Alpha-strict signature in 29 sequences of 14 unique variations (Table 2), while the rest were found in the B.1.177, C.35 and B.1.36.29 variants. The 684-653-1221 triad was part of A.27, while the 52-67-888 triad was part of Eta. The 214-215-216 insertion was part of Alpha (EPI_ISL_1920610). Seven, one and one out of the nine dyad associations involved positions between the S1/S2, S1/S1 and S2/S2 regions, respectively (Table 2). Some of these simple structures were also observed in the unique sequence population (SI File 1, F).

**Figure 2.**
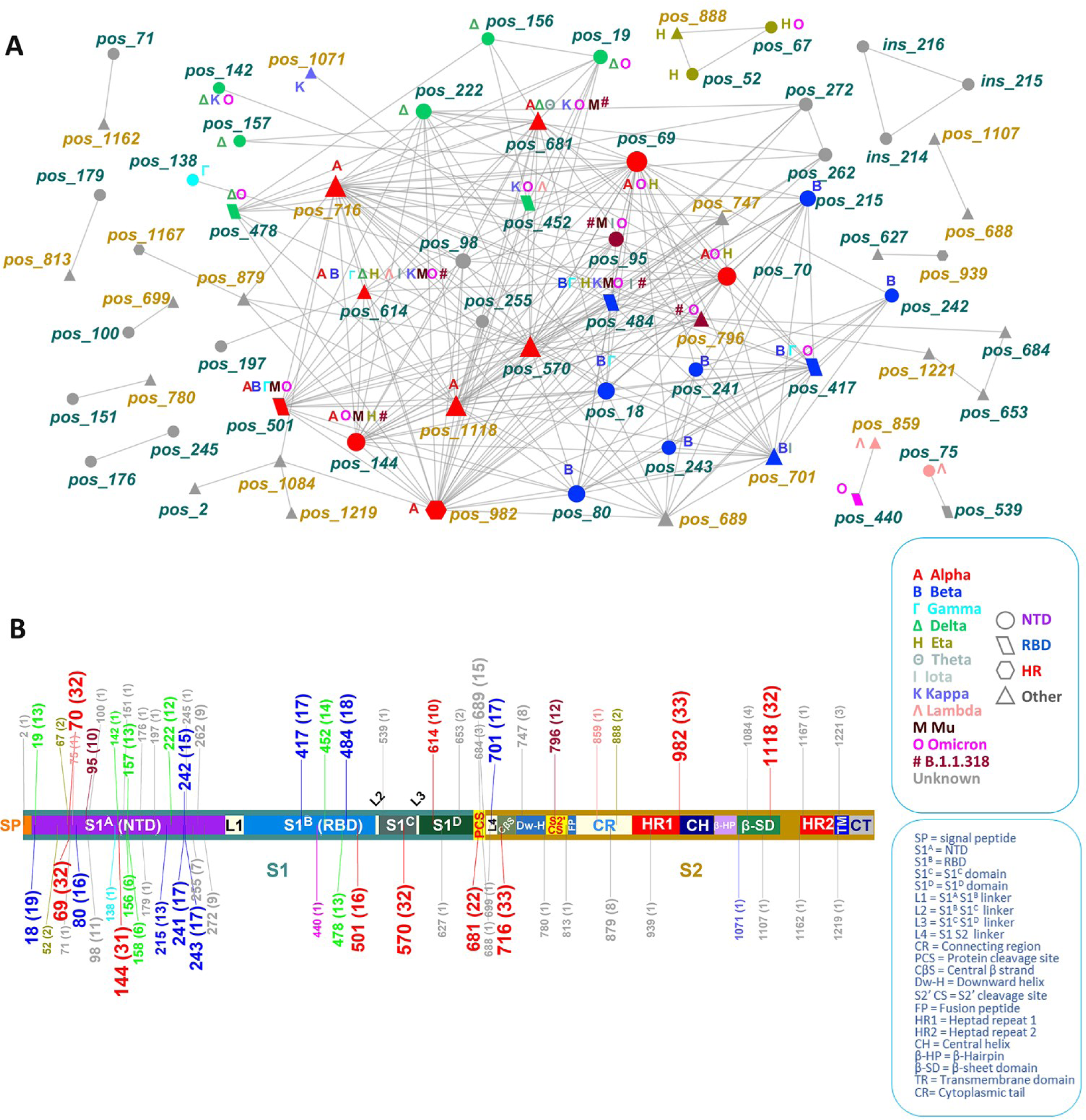
The intramolecular network **(A)** of variable positions and the corresponding spike map **(B)**. Node and map label color correspond to representative variants and shapes correspond to functional domains. Node and map label size is proportional to the frequency of observations. Numbers in labels show positions and numbers in parentheses in the map show the degree (number of connections) of each position. Text labels in nodes indicate shared positions across variants. Node label colors correspond to spike regions in the map, S1: spike region 1, S2: spike region 2, NTD: N-terminal domain; RBD: receptor-binding domain.

**Table 2.** Dyad and triad fragments found in the network. Variant, Frequency and Percentage show respectively, the variant, the number of sequences in which the network fragments were found, and the unique sequence percentage in the total sequence population (n=6,683). Ins=insertion.

| EPI | Dyads/ Triads | Variant | Region | Frequency | Percentage |
| --- | --- | --- | --- | --- | --- |
| EPI_ISL_2343867 | 627-939 | Alpha | S1-S2 | 19 | 0.28 |
| EPI_ISL_1964124 |  |  |  | 2 | 0.03 |
| EPI_ISL_2371255 |  |  |  | 1 | 0.01 |
| EPI_ISL_2233831 |  |  |  | 1 | 0.01 |
| EPI_ISL_2617350 |  |  |  | 1 | 0.01 |
| EPI_ISL_1420703 | 75-539 | B.1.177 | S1-S1 | 2 | 0.03 |
| EPI_ISL_2368640 | 71-1162 | Alpha | S1-S2 | 5 | 0.07 |
| EPI_ISL_1420662 | 100-699 | Alpha | S1-S2 | 29 | 0.43 |
| EPI_ISL_2369615 |  |  |  | 1 | 0.01 |
| EPI_ISL_2343903 |  |  |  | 1 | 0.01 |
| EPI_ISL_2369319 |  |  |  | 1 | 0.01 |
| EPI_ISL_1420684 | 179-813 | C.35 | S1-S2 | 5 | 0.07 |
| EPI_ISL_2369094 |  |  |  | 1 | 0.01 |
| EPI_ISL_2369232 |  |  |  | 1 | 0.01 |
| EPI_ISL_1497370 | 688-1107 | Alpha | S2-S2 | 1 | 0.01 |
| EPI_ISL_1420709 | 440-859 | B.1.36.29 | S1-S2 | 1 | 0.01 |
| EPI_ISL_1964143 | 176-245 | Alpha | S1-S1 | 3 | 0.04 |
| EPI_ISL_2369134 |  |  |  | 1 | 0.01 |
| EPI_ISL_2343417 | 151-780 | Alpha | S1-S2 | 3 | 0.04 |
| EPI_ISL_1731827 | 684-653-1221 | A.27 | S1-S1-S2 | 1 | 0.01 |
| EPI_ISL_1716722 | 52-67-888 | Eta | S1-S1-S2 | 1 | 0.01 |
| EPI_ISL_1920610 | Ins (214-215-216) | Alpha | S1-S1-S1 | 1 | 0.01 |

Examination of the degree distribution of the spike network indicated fitting to power law 1. (22) from a minimum degree (Xmin) of 5 [distance (Goodness of Fit, gof)=0.18, and parameter *γ*=2.01, p=0.22], but not to a Poisson distribution (Xmin=8, gof=0.023, λ=0.15, p<0.0001), implying a non-random process of network development (20, 22).

#### Spike subunit localization of network positions

The distribution of significant positions in S1 and S2 was 47 and 22, respectively. There were 39 positions with 159 links in S1, and 12 positions with 21 links in S2 (Fig, 3). Eight positions of S1 were exclusively connected to ten positions of S2 through 72 links. The S1 subnetwork mirrored the complexity of the main network in contrast to the simplistic S2 subnetwork (Fig.3). Although the difference in the number of all protein positions between S1 and S2 was about 3%, about two thirds (68%) of the subnetwork-associated positions mapped to S1. Similarly, 63% of the links were found in S1, 8% in S2 and 29% between S1 and S2. These findings indicate the distribution of epistatic events on spike, suggesting a greater, selection pressure in effect in S1, compared to S2.

**Figure 3.**
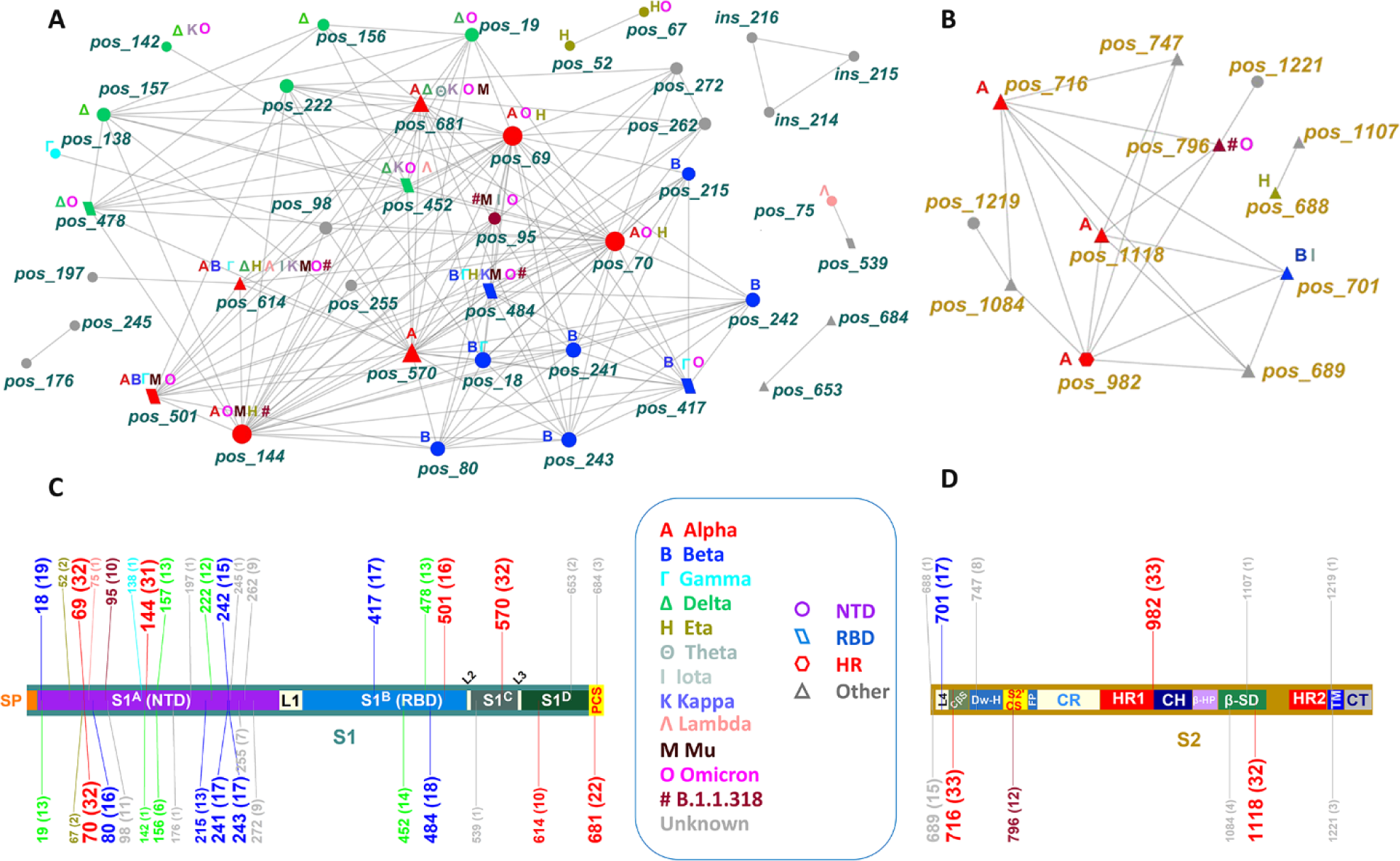
Partition of the spike network to the S1 **(A)** and S2 **(B)** regions and the corresponding mapping of network nodes in the spike protein **(C & D)**. Node color and map label color correspond to SARS-CoV-2 variants and node shapes correspond to functional domains. Node and map label size are proportional to frequency of observations. Map numbers in labels show positions and numbers in parentheses denote the network degree of each position. Text labels in nodes indicate shared positions across corresponding variants. Node label colors correspond to spike regions in the map.

#### Variant capturing in the network

The total number of positions (excluding the Alpha 214 Arg-Ala-Pro (RAP) insertion) that were associated with at least one variant and appeared in the network was 37 (out of 67, 55.2%), of which 29 was classified as at least one variant of concern (VOC), six as at least one variant of interest (VOI) and two as neither VOCs nor VOIs (B.1.1.318). The defining Alpha, Beta and B.1.1.318 sites were captured in the network in their totality (Fig.2), while the changed sites of Delta-like forms, which appeared later during evolution, were represented in the network at a percentage of 72% (8/11, site 222 included). The most connected nodes (positions 716 and 982) had degree 29 and belonged to S2, while the second most connected nodes had degree 28, with one site (570) in S1 and one (1118) in S2. Two of the S2 overconnected positions (982, 1118) were restricted in Alpha only, implying a crucial role of this region in the first wave.

#### Node connectivity in relation to VOCs and VOIs

We considered nodes fitting the power law as overconnected and examined the association between connectivity and variant type by the Pearson exact test of independence, excluding the 214-216 Alpha insertion and B.1.1.318 which was not characterized as either VOC or VOI. Of the 35 VOC/VOI-associated positions that were captured in the network, 23 of the VOC category had degree ≥5 and were thus overconnected (Table 3). Of the remaining twelve low-connectivity positions with degree <5, five (5/36) were related to at least one VOC and five (5/36) to at least one VOI. No VOI-related positions (0/36) were overconnected. The Pearson exact test of independence revealed an association between the two variables (p=0.00067, Table 3), implying a relationship between position overconnectivity in the network and variant type. Including B.1.1.318 in the analysis as either VOC or VOI, or the 214-216 insertion as VOC (Alpha)-related, did not affect considerably the significance of the test.

**Table 3.** Characteristics of the 37 VOC- and VOI-related sites captured in the network.

|  | Position | Degree | Region | Domain | Function | Variant |
| --- | --- | --- | --- | --- | --- | --- |
| 1 | 18 | 16 | S1 | S1<sup>A</sup>(NTD) | NAb | B, Γ |
| 2 | 19 | 8 | S1 | S1<sup>A</sup>(NTD) | NAb | Δ, O |
| 3 | 52 | 2 | S1 | S1<sup>A</sup>(NTD) | NAb | H,O |
| 4 | 67 | 2 | S1 | S1<sup>A</sup>(NTD) | NAb | H |
| 5 | 69 | 26 | S1 | S1<sup>A</sup>(NTD) | NAb | A, O, H |
| 6 | 70 | 20 | S1 | S1<sup>A</sup>(NTD) | NAb | A, O, H |
| 7 | 75 | 1 | S1 | S1<sup>A</sup>(NTD) | NAb | Λ |
| 8 | 80 | 16 | S1 | S1<sup>A</sup>(NTD) | NAb | B |
| 9 | 95 | 10 | S1 | S1<sup>A</sup>(NTD) | NAb | B.1.1.318, I, K |
| 10 | 138 | 1 | S1 | S1<sup>A</sup>(NTD) | NAb | Γ |
| 11 | 142 | 1 | S1 | S1<sup>A</sup>(NTD) | NAb | Δ, O |
| 12 | 144 | 19 | S1 | S1<sup>A</sup>(NTD) | NAb | A, H, M, O, B.1.1.318 |
| 13 | 156 | 6 | S1 | S1<sup>A</sup>(NTD) | NAb | Δ, Λ |
| 14 | 157 | 2 | S1 | S1<sup>A</sup>(NTD) | NAb | Δ |
| 15 | 215 | 13 | S1 | S1<sup>A</sup>(NTD) | NAb | B |
| 16 | 222 | 12 | S1 | S1<sup>A</sup>(NTD) | NAb | Δ, B.1.177 |
| 17 | 241 | 8 | S1 | S1<sup>A</sup>(NTD) | NAb | B |
| 18 | 242 | 7 | S1 | S1<sup>A</sup>(NTD) | NAb | B |
| 19 | 243 | 8 | S1 | S1<sup>A</sup>(NTD) | NAb | B |
| 20 | 417 | 17 | S1 | S1<sup>B</sup>(RBD) | ACE2 | B, Γ, O |
| 21 | 440 | 1 | S1 | S1<sup>B</sup>(RBD) | ACE2 | O |
| 22 | 452 | 10 | S1 | S1<sup>B</sup>(RBD) | ACE2 | Δ, K, Λ, E, O |
| 23 | 478 | 9 | S1 | S1<sup>B</sup>(RBD) | ACE2 | Δ, O |
| 24 | 484 | 14 | S1 | S1<sup>B</sup>(RBD) | ACE2 | B, Γ, K, M, Z, H, Θ, I, B.1.1.318, O |
| 25 | 501 | 16 | S1 | S1<sup>B</sup>(RBD) | ACE2 | A, B, Γ, Θ, M, O |
| 26 | 570 | 28 | S1 | S1<sup>C</sup> domain | Unknown | A |
| 27 | 614 | 10 | S1 | S1<sup>D</sup> domain | Unknown | A, B, Γ, Δ, E, Z, H, Θ, I, K, Λ, M, B.1.1.318, O |
| 28 | 681 | 19 | S1 | PCS | Cleavage | A, Δ, Θ, K, M, B.1.318, O |
| 29 | 701 | 17 | S2 | L4 | Unknown | B, I |
| 30 | 716 | 29 | S2 | CβS | Unknown | A |
| 31 | 796 | 12 | S2 | S2' | Cleavage | B.1.1.318, O |
| 32 | 859 | 1 | S2 | CR | Unknown | Z, Λ |
| 33 | 888 | 2 | S2 | CR | Unknown | H |
| 34 | 982 | 29 | S2 | HR1 | Fusion | A |
| 35 | 1071 | 1 | S2 | β-SD | Unknown | K |
| 36 | 1118 | 28 | S2 | β-SD | Unknown | A |

ACE2: Angiotensin-converting enzyme 2 (ACE2) receptor-binding; NAb: Neutralizing antibody binding; RBD: Receptor binding domain; HR1: heptad repeat 1; PCS: protease cleavage site; L4=S1-S2 linker, CβS= central beta strand; S2': cleavage site; L5: fusion peptide-HR1 connecting region; β-SD: β-strand domain.

#### Node association to Time

As the varying sequences were distributed over the three pandemic waves, we examined the potential association of variable positions to time, adding the dates of sample collection as levels of the “Date” time-related factor. We found highly significant associations of some positions to Date (SI Fig.3), suggesting a rapid adaptation rate (Table 4). Overall, 26/170 positions were related to time, of which 25 were included in the network and one (site 210) was excluded having lowest exact p=0.002 and highest Z-score =0.*214*. *Over*connectivity (degree ≥*5*) characterized 19 of the time-related positions, low connectivity (0<degree<5) characterized four positions, and no connectivity (degree=0) characterized the remaining three positions. Twenty-three of these positions have been reported in the mutational constellation of at least one variant, while one position (number 879) has not been reported for any variant. These results suggest a dynamic time-dependent process of aminoacidic changes among variant positions, captured by the network.

**Table 4.** Time-related variable positions of the spike protein (n=1,078, p≤0.001, Pearson exact test). Time shows significances at the Day level.

| No. | Spike Position | Degree | Related Variant(s) |
| --- | --- | --- | --- |
| 1 | 18 | 20 | B, $\Gamma$ |
| 2 | 19 | 13 | $\Delta$, O |
| 3 | 69 | 32 | A, O, H |
| 4 | 70 | 32 | A, O, H |
| 5 | 144 | 32 | A, H, M, O, B.1.1.318 |
| 6 | 156 | 6 | $\Delta$, $\Lambda$ |
| 7 | 157 | 13 | $\Delta$ |
| 8 | 158 | 6 | $\Delta$ |
| 9 | 210 | 0 | Omicron (BA.2.75) |
| 10 | 222 | 12 | $\Delta$ |
| 11 | 215 | 13 | B, B.1.616 |
| 12 | 241 | 17 | B |
| 13 | 243 | 17 | B |
| 14 | 417 | 17 | B, $\Gamma$, O |
| 15 | 452 | 14 | A, $\Delta$, K, $\Lambda$, O |
| 16 | 478 | 13 | $\Delta$, O |
| 17 | 570 | 32 | A |
| 18 | 614 | 10 | A, B, $\Gamma$, $\Delta$, E, Z, H, $\Theta$, I, K, $\Lambda$, M, B.1.1.318, O |
| 19 | 681 | 22 | A, $\Gamma$, $\Delta$, $\Theta$, K, M, B.1.1.318, O |
| 20 | 701 | 17 | B, I |
| 21 | 716 | 33 | A |
| 22 | 879 | 8 | Undefined |
| 23 | 982 | 33 | A |
| 24 | 1073 | 0 | Undefined |
| 25 | 1104 | 0 | Undefined |
| 26 | 1118 | 32 | A |

#### Network evolution with time

We further examined the evolution of associated spike positions over time, partitioning the dataset by month and creating the corresponding networks. We found no significant associations between spike positions before 2020-12-01. The total number of sequences up to that date was 135, with 25 variable positions. A cohesive module of nine Alpha-related positions along with dyad 18/222 appeared in December 2021, forming 11 network nodes and 37 links (Fig.4A). In January 2021, dyad 18/222 along with some extra positions (262/272) were linked to the Alpha-related network structure (Fig.4B). Meanwhile, a group of eight tightly connected Beta-related nodes and four dyads appeared independently of other network nodes, increasing its size to 29 nodes and 77 links. In February 2021, some positions of the Alpha- and Beta-related structures started becoming connected and dyad 98/747 joined the main network which amounted to 33 nodes and 117 links (Fig.4C). In March 2021, some new low degree nodes were linked in the network periphery, along with dyad 879/1167, while a group of tightly connected Delta-related positions appeared independently, resulting in a further increase to the network size (63 nodes/234 links, Fig.4D). This structure was the direct precursor of the final network of 69 nodes and 252 links (Fig.2).

**Figure 4.**
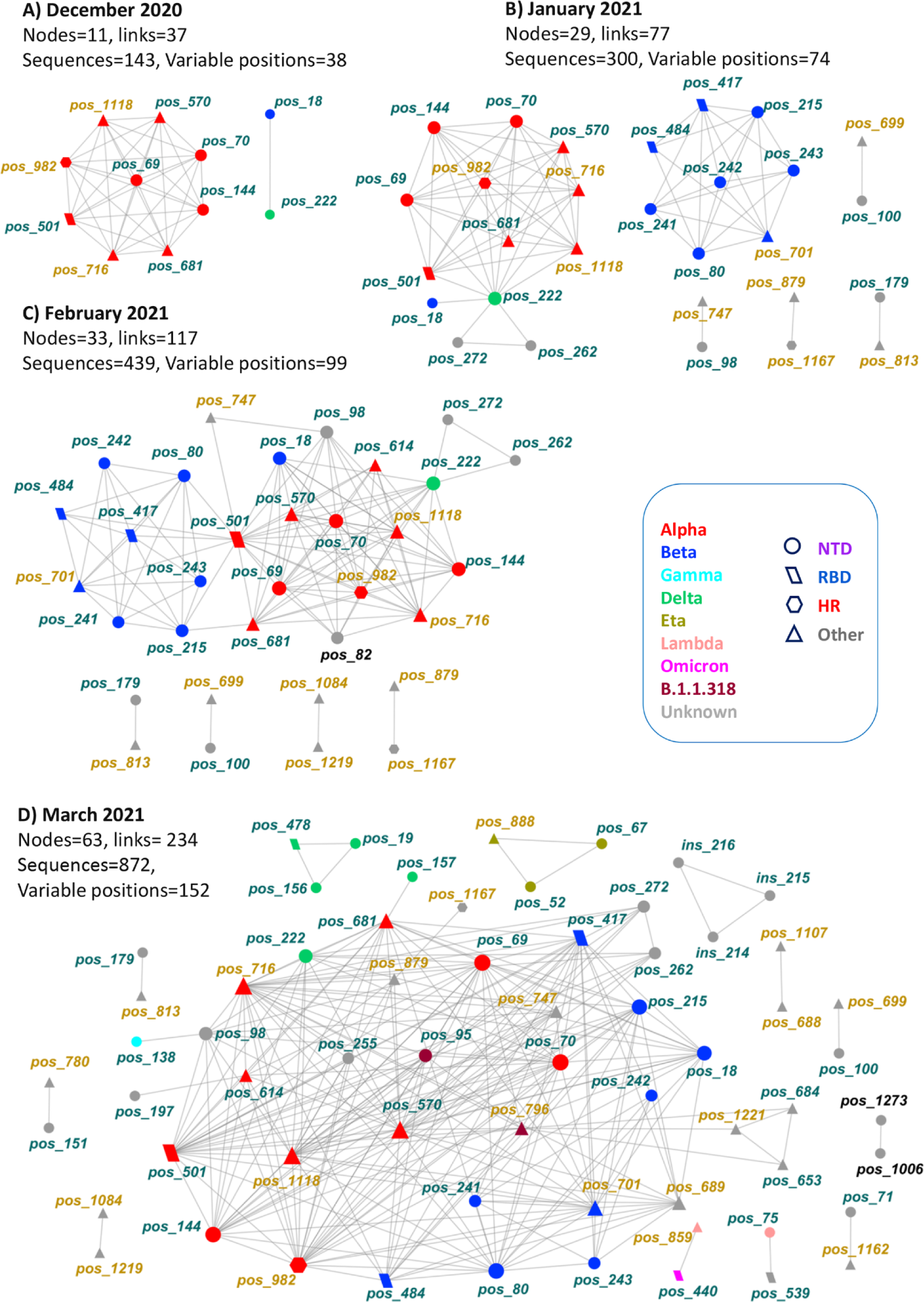
Evolution of the spike intramolecular network over time (December 2020 - March 2021). Node color indicates variant or spike subunit location (S1, dark green; S2, light brown), with black indicating nodes missing from the final network. Node size in each graph is proportional to degree.

During this dynamic process, position 82 with eight connections appeared in intermediate steps, but later lost its links and was absent from the final network. Position 18, a known occasionally reported Beta site, appeared independently of Beta-related positions in the earliest network form linked to position 222. However, position 18 gained more links subsequently and was tightly connected to Beta-related positions in the final steps. The first occurrence of dyad 18- 222 in the population was recorded on 2020-10-12 (EPI_ISL_2302048) and the first appearance of Beta was recorded on 2021-01-14 (EPI_ISL_1716714), implying an affinity between the two positions.

#### Relationship of cliques and communities to variants

Networks hold structural information in the form of cliques of connected nodes and communities characterized by interconnected *k-*cliques (*k≥*3) via *k-*1 nodes (23). In intramolecular viral protein networks, cliques may represent evolving relationships over time among nodes with shared properties that ensure the community’s functionality, thus allowing for predictions on the properties of unknown nodes (23). We further explored network modularity in terms of *k-*cliques and functional communities and found 25 cliques (SI File 2) with *k* ranging between three and 11. The maximum number of communities was four, observed at the *k*-3 and the *k*-7 clique resolutions,. The four *k-7* communities had 7, 7, 22, and 8 nodes, respectively, were numbered consecutively from one to four and studied further. Based on their mutational profiles, the first second and fourth communities approximated the Alpha, Beta and Delta variants. Thus, the Alpha-like community included the Alpha known positions 570, 716, 982, 1118, shared with Beta and Delta communities, while position 222 was shared with the Delta community and two unknown positions (262, 272) later found as Alpha-related (24). The Beta-like community included the known Beta deletion 241-243, the Beta positions 80, 417, 701, and the unknown position 689. These four positions were shared with the third community. The third, largest community, included all B.1.1.318 positions (e.g., 95, 144, 484, 614, 681, 796) as well as multifunctional sites of other variants and two unknown positions, namely 98, and 747. The Delta-like community included the known Delta positions 19, 452, 478, as well as positions 570, 681, 716, 982 and 1182. Position 681 was shared with the Beta-like community and all the rest were shared with the Alpha-like community.

These findings suggest excessive interactions among variable positions in the course of virus evolution, depicted as a pooled community of heterogeneous virus forms, linked to more well characterized communities (e.g. Alpha, Beta) or the under development Delta community. The overlapping positions could imply important multifunctional roles for the sharing variants at given momenta of virus evolution and suggest epistatic interactions between positions, later known to characterize specific VOCs. For example, Delta positions 19, 452 and 478, belonging to the same community as Alpha positions, were found in relation to Alpha variants in real-world populations (e.g., EPI_ISL_2370539, EPI_ISL_2343708 and EPI_ISL_2364971, respectively).

#### From network cliques and communities to significant sequences

We also examined the occurrence of clique positions in real-world sequences, using three positions (a 3-*k* sub clique) of the *4-k* and 8-*k* cliques (SI File 3) as a paradigm, involved in the formation of the Delta-related community (Fig.5). Theoretically, the total number of sequences containing all possible positional combinations of a clique is the product of the number of amino acids in each position. In the studied 3-*k* clique there were three positions (19, 452, 478) each having two amino acids (including deletions and unchanged residues). The number of sequences holding all possible combinations was eight (SI File 3). The most frequently detected combinations were “19R, 452R, 478K”, and “19T, 452L, 478T” scoring seven, and three, counts, respectively. The unmutated positions had the form “T19, L452, T478” and scored 1068 counts. None of the rest five potential combinations was present in our sample (SI File 1) since it probably led to less fit variants or deleterious mixes of mutations in those early stages of virus evolution.

**Figure 5.**
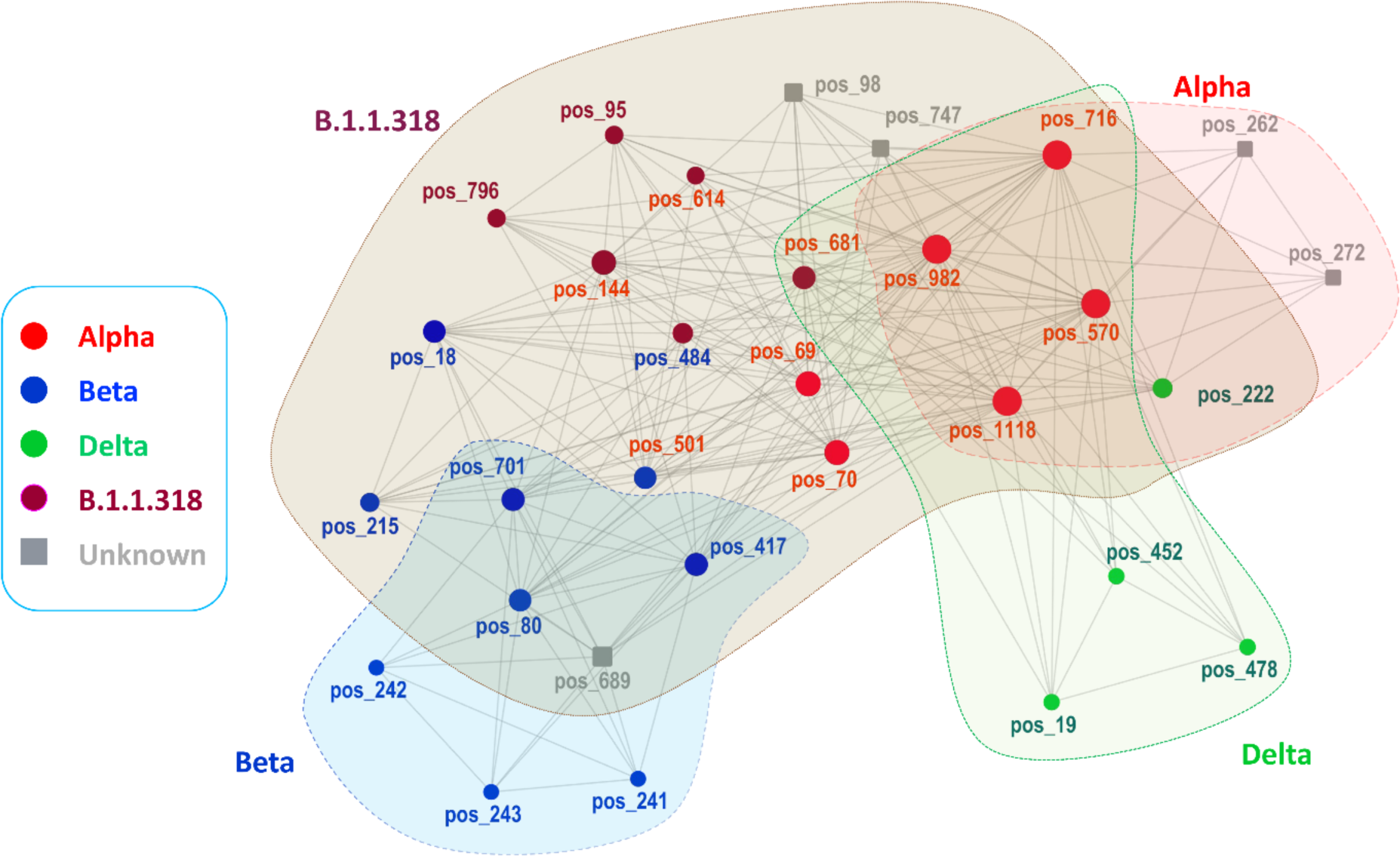
The four *k*-7 clique communities of the intramolecular network of the SARS-CoV-2 spike protein. Node size is proportional to node degree. Round nodes show positions reported in at least one variant and square nodes show predicted positions, not reported (i.e., 98, 747, 689) in any variant, or reported in Alpha variant (i.e., 262, 272) at the end of the study period (April 26, 2021). Red, blue, green and brown nodes correspond to positions in the Alpha, Beta, Delta and B.1.1.318 variants. When in different communities, the color of nodes and labels corresponds to more related communities/variants and original variants, respectively. Grey color denotes positions not reported in any variant by the end of study period.

## DISCUSSION

We examined the distribution of all SARS-CoV-2 sequences available in GISAID from Greece, covering the first two and most of the third pandemic waves. Probing *in silico* by strict variant signatures we determined the circulating variants which comprised four VOCs (Alpha, Beta, Gamma and Delta-like), two VOIs (Eta and Kappa) and one undefined (B.1.1.318) lineage. Exploration of the aminoacidic differences of each sequence from the prototypic further unveiled 652 unique variations of the initial sequence, with the number of mutated positions ranging between one and 15. The frequencies and the distribution of these unique sequences varied considerably, reflecting the potential of the virus population for adaptation.

A common feature of the evolving sequences was the gradual growth of the number of changed positions. This temporal increase suggests an evolutionary process likely controlled by epistatic phenomena, in which the change of an amino acid in one position allosterically affects others (25, 26, 7). Virus evolution is known to be affected by epistasis, which promotes trajectories conditioned on already occurred mutational events (27).

The contingency tables approach that we used requires the assumptions of random sampling and independence among samples, which may be violated in some cases, e.g. from the potential clustering of samples due to the tree-like nature of genomic co-variational data. To eliminate possible phylogenetic bias and random noise between pairs of positions, we calculated the mutual information score applying APC (28) and the corresponding Z-scores,. Since positions having Z-scores ≥ 2 were shown to be associated in an eukaryotic protein (28), we used Z-scores ≥ 2 coupled with exact p ≤ 0.001 cutoffs to analyze further our spike associations.

The discovered associations between positions and between positions and time, support a time-dependent epistatic mechanism in virus evolution. Most of these positions belonged to known VOCs and VOIs, while others belonged to unassigned variants. The revealed associations were overlapping, with some shared positions resembling complex systems. Complex epistatic effects on spike functionality has been suggested (7), while complex networks of associated positions have been observed in viral (6, 29, 30) or other proteins (31).

We applied network analytical tools to explore mutational dynamics, important positions and structural relationships. Our analysis revealed a temporally evolving network, growing in size and complexity with time. The ordered network path that includes all nodes in ascending order could represent a single theoretical spike sequence, comprising all candidate mutational sites and associations. Across evolution, only a handful of these variable positions could be engaged, most likely affected by initial conditions (27), resulting in unique aminoacidic combinations.

Already mature variants, entering new populations *per se* (e.g., Alpha or Beta) may also engage mutations at positions beyond their strict signatures as a result of microenvironmental adaptations, that could be depicted in a network. The observed mutational Alpha combinations, beyond its strict signature that engaged dyads present in the network (e.g. 627-939, 75-539), may indicate adaptation paths after Alpha variant introduction in the region. Thus, the presented network could reveal complex relationships among variable positions in interrelated sites and unveil the evolutionary dynamics of VOCs and VOIs formation in relation to their mutational profiles.

Network development also reflected the mutational status of real-world sequences at elapsing time spans. As mutations in novel spike positions were gradually adopted over time, new variable sites were added, increasing the number of significant associations. Associations among variable positions are gained by mutational adaptations during virus evolution. These may be revealed as dyads, which also represent 2*k*-cliques that are the structural blocks of the entire network, mirroring instantaneous epistatic associations in the virus population. Such dyads may be independent, or they may relate to other positions, forming triads or higher order structural blocks. Interestingly, this process captures predictive information as, two or more positions associated to another, have a higher probability to relate to each other (21). Such information could be exploited through relevant network centralities such as clustering coefficient, eigenvector, betweenness and other centralities. Thus, a growing intramolecular network, as shown in our work, could depict the complexity of aminoacidic associations among virus isolates, providing information on the dynamics of fluctuating protein conformations.

During the first wave, we found no significant position associations, possibly due to the few amino acid changes at the early pandemic stages, apart from the crucial D614G substitution (12, 13). Another reason could be the small number of available sequences then. Significant associations appeared in the network towards the end of the second wave, uncovering all strictly- Alpha signature positions and the 18/222 dyad. With time, new positions of known VOCs and VOIs as well as positions unassigned to any variant became evident, reflecting the appearance of novel aminoacidic combinations and the momentary formation of lineages of unknown epidemiological role.

The node degree distribution fitted partially a power law, supporting a gradual adaptation hypothesis. The network development initially involved the formation of simple dyads, many of which progressively merged wtih peripheric nodes, suggesting constrained epistatic expansion (27), showcasing the complexity (29, 5) of aminoacidic interactions (7). Progressively, some positions gained more associations than others, implying a potential biological role, further supported by the findings of significant associations between overconnected positions and VOCs that provided a direct relation of network degree to virus functionality and adaptation.

In support of this inference, the final-step network captured all or almost all positions of circulating variants and revealed numerous fragmented dyads and triads, representative of mutating sites of evolving sequences in the population. These structures were found in various forms, such as mature (e.g. Alpha) and undefined variants, the latter dominating during the early pandemic phases. Most of these fragments involved paired associations between positions of the S1 and the S2 regions. As these regions display distinct functionality, mainly specific for antigenic escape and membrane fusion, a positive selection mechanism concurrently targeting immunogenetic drift and particle fusion cannot be ruled out, particularly at the early stages of virus evolution or during the entry of mature variants in new regions.

To reveal potential patterns related to real-world sequences, we further analyzed the structure of the final-step network that represented the cumulative mutational associations between positions up to the end of April 2021. We found 25 cliques, which at different resolutions (number of clique nodes) formed one to four communities. Cliques and communities are hierarchical structures in networks, which could be exploited to gain information on node properties (23), such as protein conformation, functionality, and allosteric communication (32). For example, the fourth clique of the network captured the instantaneous relationship of positional associations of the Delta variant. The dynamics of these associations were depicted in the Delta-like community, which likely hold relevant functional and structural information. Indeed, positional associations in this community matched well with a recently reported structural representation of the Delta variant (32). Thus, position 19 in the NTD, as well as positions 452 and 478 in the RBD and 681 in the furin cleavage site are structurally associated and they are also tightly connected, comprising, the first three, the unique sites of the Delta-like community and the fourth an overlap with the Alpha, Delta and B.1.1.381 communities. Overlapping nodes between communities may signify epistatic interactions among different variants or virus forms in the population towards structural conformations of new variant formation.

Unknown nodes found inside a functional community are likely to share properties with the rest of the community nodes. Increasing or decreasing the number of clique nodes (*k*) resembles zooming in and out of a microscope’s field, providing more or less detailed information on nodes of interest (23).

Analysis of the network at the 7-*k* resolution revealed four overlapping communities capturing positions of the Alpha, Beta, Delta and B.1.1.318 variants. Three positions, namely 98, 747, 689 of the large (third) community, and positions 262 and 272 of the Alpha community, have not been assigned to any variant. Position 98 was initially associated with site 747 and became progressively connected to B.1.1.318, Beta and Alpha, implying a continuous overlapping with sites of these variants over time. Similarly position 689, located proximally to the important S1/S2 cleavage site, was an extra Beta mutating site shared with the third community. Positions 262 and 272 were found early in the pandemic in many unassigned variants, closely associated to position 222 which has been reported in some Delta lineages. Recently, positions 262 and 272 were found in association to each other, while positions 262 has been reported in connection to the Alpha variant in phylogenetic analyses, confirming the prediction of its allocation within the Alpha community (24).

Two types of significant aminoacidic associations in pairs of variable positions could be defined, involving either mutual compatibility or mutual exclusion. The former derives from adaptation between two variable positions in the same sequence and the latter concerns aminoacidic changes of two positions in different sequences. We used as an example a 3-*k* clique of the Delta-associated community to find the mutational profile of sequences in our sample. We constrained our approach to only three positions to keep explored sequences at a manageable size. Theoretically, the total number of sequences carrying all possible combinations, including the unmutated sequence, was eight. However, only two sequences were found to hold different mutational combinations. Most sequences carried four aminoacidic adaptations, thus showing mutual compatibility. The less frequent sequences had one and three changes in these positions, showing mutual exclusion or mixed dependencies, respectively. Hence, cliques and communities of interest could be exploited programmatically to uncover significant aminoacidic combinations in terms of antigenicity and other biological properties in real-world sequences.

In summary, based on noise-free statistically significant associations between SARS-CoV- 2 spike variable positions, we developed an analytic approach that offers a novel way for understanding virus epidemiology and evolution by linking network structural aspects to mutational patterns in the spike sequence population.

## Materials and Methods

### Used software

Dataset preparation and network analysis were performed in “R” (33), except otherwise stated, by using or programming functions of base “R” and the following libraries: “dplyr” (34), “tidyr” (35), “GISAIDR” (36), “ggplot2” (37), “ggrepel” (38), “openxlsx” (39), “msa” (40), “poweRlaw” (41), “igraph” (42), “RCy3” (43), “biostrings” (44) and “Bios2cor” (45).

### Multiple sequence alignment

Whole SARS-CoV-2 genomes available from Greece for the period February 29, 2020 - April 26, 2021 were retrieved from GISAID (46) (accessed on 2021- 05-01) and used for network analysis. Sequences were aligned using ClustalOmega (47) with default parameters via the “msa” “R” library. Sequences >25,120 nucleotides, or with >10 ambiguous characters, were discarded. The spike gene was isolated from the alignment and translated to the corresponding amino acids using the “biostrings” R library. Spike-coding sites were mapped to the alignment from the index virus (Wuhan-Hu-1) sequence (GenBank Accession number NC_045512.2) excluding stop codons. Insertions and deletions were annotated accordingly, using “R” scripts, to keep the prototypic sequence numbering. Conserved sites were excluded. The resulting alignment, containing annotated variable positions, was used for pairwise analysis.

GISAID records containing spike protein sequences were also retrieved (GISAID accessed on 2021-06-01) and used for mutational analysis. Sequences were aligned using the MAFFT plugin of the Geneious software (Geneious Prime 2019.2.3, Biomatters Ltd, Aukland, New Zealand). Sequences with more than five ambiguities were removed and conserved positions were excluded. The remaining sequences were annotated and renumbered as described above. GISAID metadata that included sample information on collection dates and location, were also retrieved.

### Pairwise associations of amino acid positions in spike

Pairwise position associations in the spike protein of the whole genome dataset were examined in R × C (r=rows, c=columns) contingency tables by the Pearson Chi-Square exact test of independence, assuming a multinomial joint distribution for the cell scores (48, 49). The corresponding *p* exact values were obtained for each table of paired positions using the CROSSTABS algorithm of the SPSS program by looping sequentially all pairwise position combinations. The resulting values were retrieved by XPATH scripts of the XML language (50). The procedure was implemented by code written in Visual Basic (Visual Studio Express 2015, Microsoft corp., Redmond, WA, USA) using the IBM SPSS Statistics – Integration Plug-in for Microsoft.NET (51) to access the SPSS engine (IBM SPSS Statistics for Windows, Version 23.0. Armonk, NY: IBM Corp.).

In each table, rows and columns represented the levels (amino acids) of examined positions and cells denoted the frequencies of amino acid combinations. The number of pairwise

combinations was given by the formula, 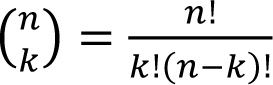 where *n* is the total number of items and *k* is the number of chosen items without replacement. In the case of the “Time” variable, rows denoted the time levels of dates and columns denoted the levels of each amino acid position. For a given R × C contingency table, the Pearson exact test examines the null hypothesis *H*_0_ : π*_i, j_* = π*_i+_*, π*_i+j_* for all cells (*i, j)* against the alternative hypothesis *H*_1_ : π*_ij_* ≠ π*_i+_*, π*_i+j_*, where π*_i+_* = 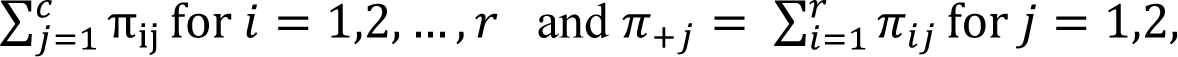 (8).

Exact algorithms calculate the probability for each R × C table by summing up the non-parametrically-obtained (exact) probabilities (48, 8) of all contingency tables, which, under the null hypothesis, are less likely to occur than the table examined within a sample space defined by all R × C tables that have the same marginal sums as the examined one (48, 9, 52, 8). As exact calculations are computationally intensive, the exact values were approximated with extremely high accuracy (52) by the Monte Carlo algorithm in the SPSS program, using a random sampling of 10,000 tables for each pair. The employment of exact tests was necessary to minimize Type I error (52) and overcome potential problems of imbalanced and sparse contingency tables for which asymptotic tests are often inaccurate (10, 49).

### Phylogenetic signal, random noise and recombination tests

Associations between positions might include random noise and phylogenetic information (53, 28). Several methods have been developed to avoid (54, 55) or exclude such information on paired associations. We used the average product correction (APC), via the “Bios2cor” library in “R”, that eliminates phylogenetic and random background signals (28). To test the possibility of recombination events that could interfere with paired associations, we tested the protein-coding nucleotide alignments of the spike gene, using relevant algorithms of the RDP5 software (56).

### Construction, visualization, and analysis of the intramolecular spike network

The subpopulation of spike position pairs with probability values ≤0.001 in the Pearson exact test and Z-score ≥ 2 was used for network construction.

The network of position associations was created by programming functions of the “igraph” library in “R” and displayed in Cytoscape (v.18.2) (57) using the “R” “RCy3” library of the “Bioconductor” environment. Functional communities in the network were identified using the clique percolation algorithm through the “cfinder” software (23). Fitting the network degree distribution to power law and Poisson distributions was based on maximum likelihood parameter estimation and on bootstrap uncertainty estimation (58, 41, 59), both implemented by programming the “poweRlaw” library in “R”, allowing 5,000 bootstraps. Fitting to power law was verified by employing appropriate functions of the “R” “igraph” library. The association of the network degree to the frequency of adapted aminoacidic changes in each position was assessed by regression analysis and by the non-parametric test of Spearman *ρ*, using “R”.

### Mutational profiles

To assess mutational profiles, we aligned all Greek spike protein sequences available in GISAID for that period [1,078 translated whole genomic+5,605 partial (spike-only) = 6,683 sequences] and probed for variant signatures by programming functions of “R” libraries. Aminoacidic changes were assessed by comparing the number and type of residues in the corresponding positions. We used local polynomial regression (loess) and linear regression in “ggplot2” library in “R” to obtain trend lines of the number of aminoacidic changes over time. The “lm” function in “R” was used for parameter estimation of linear regression.

## Data availability

The data that support the findings of this study are available upon request.

---

**SI Figure 1.**
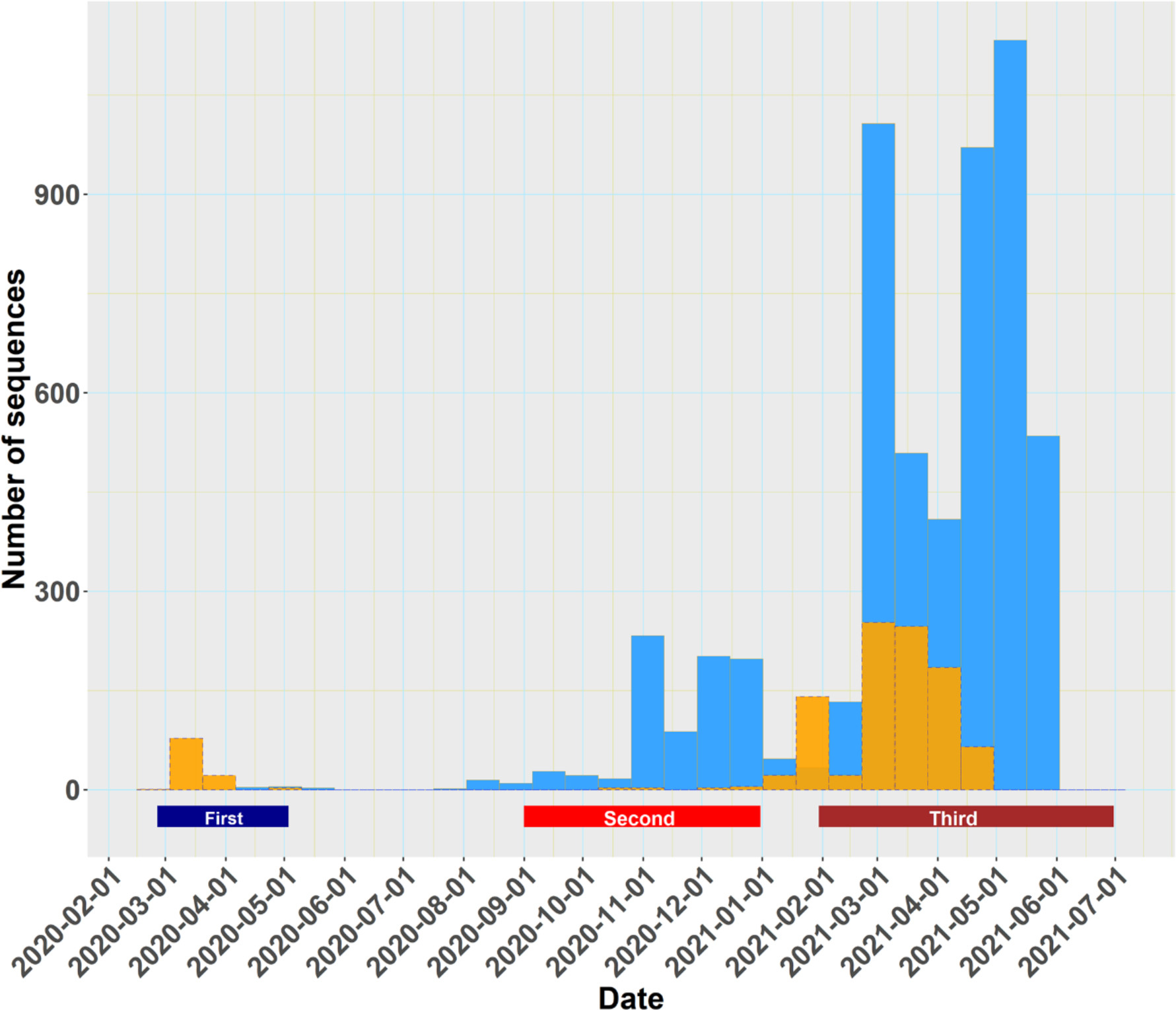
Distribution in time of whole genome (orange) and partial genome (spike only, cyan) sequences used for network construction and mutational frequency analysis. Labels indicate the first, second and third pandemic waves.

**SI Figure 2.**
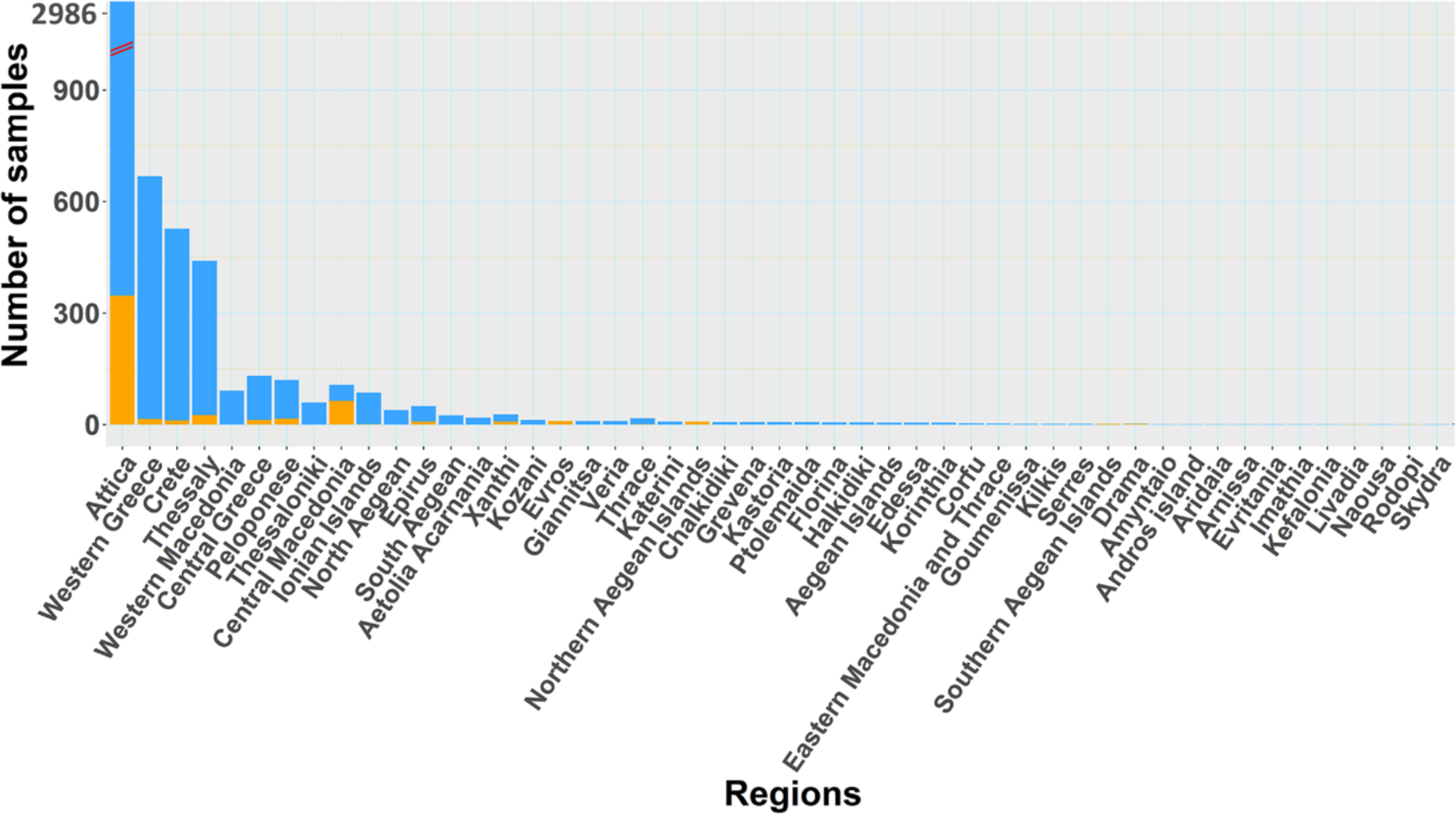
Stacked bar plot showing regional distribution of whole genome (orange) and partial genome (spike only, cyan) sequences used for network construction and mutational frequency analysis.

**SI Figure 3.**
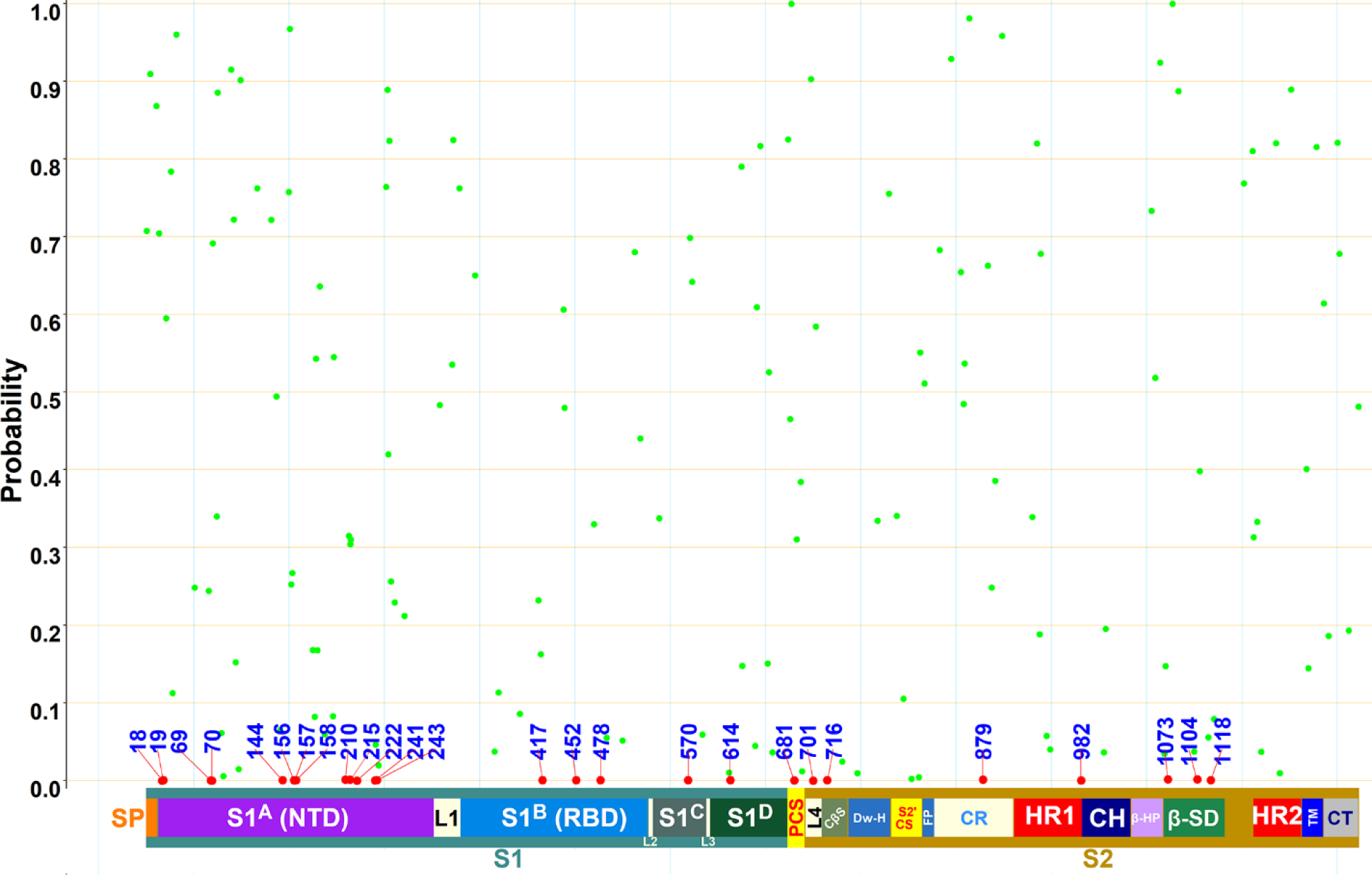
Probabilities of identified variable spike protein positions (n=170) related to time. Green dots indicate probability values >0.001 of the Day variable. Numbers with dots indicate positions. Red color indicates significances (p≤0.001) for the corresponding positions. The map represents the spike protein. The dark green and brown bars display the S1 and S2 regions, respectively. SP= signal peptide, NTD=N’ terminal domain, RBD = Receptor binding domain, HR=Heptad Repeat (1 and 2).

**SI Table 1.**
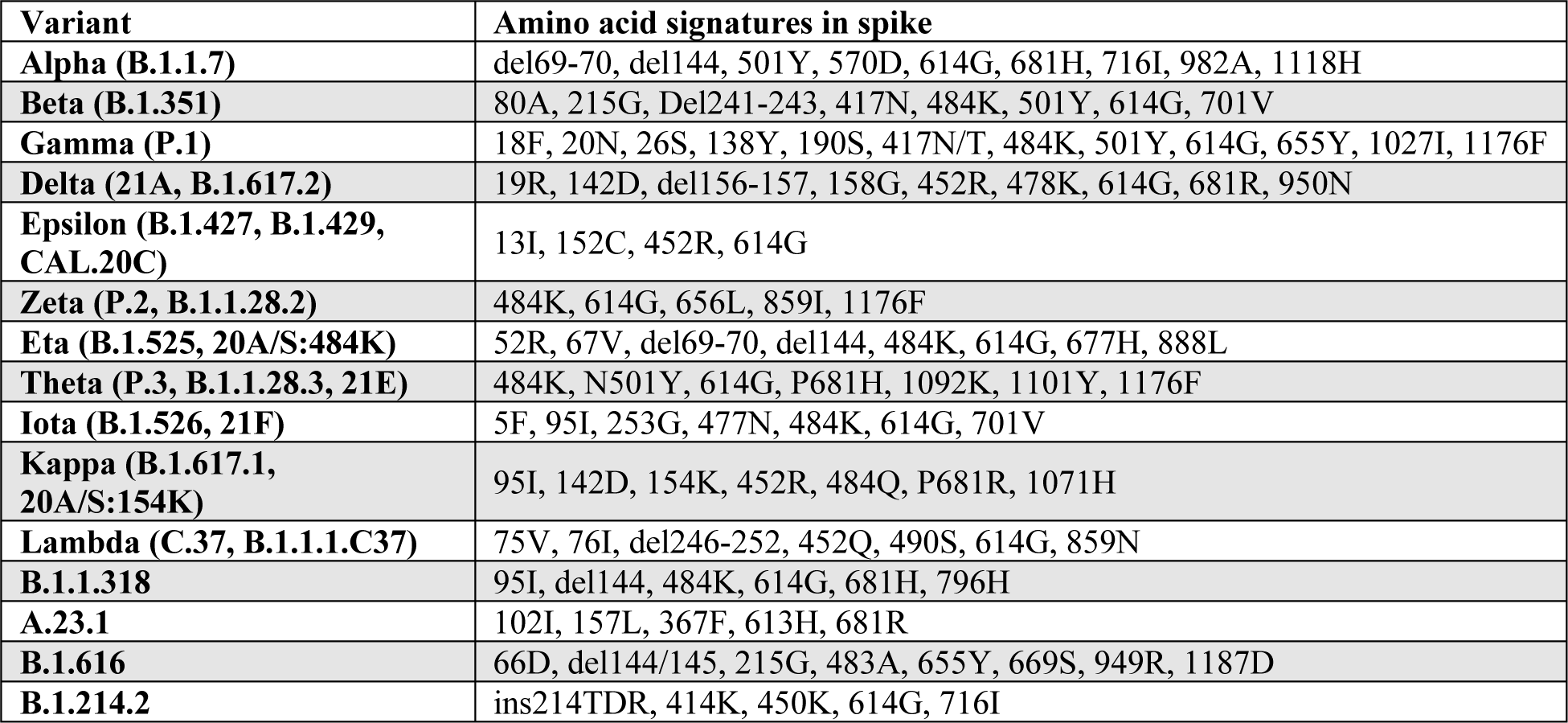
Selected SARS-CoV-2 variant-defining spike amino acid signatures.

| **Variant** | **Amino acid signatures in spike** |
| --- | --- |
| **Alpha (B.1.1.7)** | del69-70, del144, 501Y, 570D, 614G, 681H, 716I, 982A, 1118H |
| **Beta (B.1.351)** | 80A, 215G, Del241-243, 417N, 484K, 501Y, 614G, 701V |
| **Gamma (P.1)** | 18F, 20N, 26S, 138Y, 190S, 417N/T, 484K, 501Y, 614G, 655Y, 1027I, 1176F |
| **Delta (21A, B.1.617.2)** | 19R, 142D, del156-157, 158G, 452R, 478K, 614G, 681R, 950N |
| **Epsilon (B.1.427, B.1.429, CAL.20C)** | 13I, 152C, 452R, 614G |
| **Zeta (P.2, B.1.1.28.2)** | 484K, 614G, 656L, 859I, 1176F |
| **Eta (B.1.525, 20A/S:484K)** | 52R, 67V, del69-70, del144, 484K, 614G, 677H, 888L |
| **Theta (P.3, B.1.1.28.3, 21E)** | 484K, N501Y, 614G, P681H, 1092K, 1101Y, 1176F |
| **Iota (B.1.526, 21F)** | 5F, 95I, 253G, 477N, 484K, 614G, 701V |
| **Kappa (B.1.617.1, 20A/S:154K)** | 95I, 142D, 154K, 452R, 484Q, P681R, 1071H |
| **Lambda (C.37, B.1.1.1.C37)** | 75V, 76I, del246-252, 452Q, 490S, 614G, 859N |
| **B.1.1.318** | 95I, del144, 484K, 614G, 681H, 796H |
| **A.23.1** | 102I, 157L, 367F, 613H, 681R |
| **B.1.616** | 66D, del144/145, 215G, 483A, 655Y, 669S, 949R, 1187D |
| **B.1.214.2** | ins214TDR, 414K, 450K, 614G, 716I |

**SI Table 2.** Dates of first reports of SARS-CoV-2 index virus and variants in Greece.

| **Variant** | **GISAID**<br>**accession number** | **Year** | **Month-Day** |
| --- | --- | --- | --- |
| Wuhan-Hu-1 | EPI_ISL_2232041 | 2020 | Feb 26 |
| Alpha | EPI_ISL_2301679 |  | Dec 11 |
| Beta | EPI_ISL_1716721 | 2021 | Jan 14 |
| Gamma | EPI_ISL_2454299 |  | May 14 |
| Delta-like* | EPI_ISL_1716736 |  | Mar 23 |
| Eta | EPI_ISL_1716722 |  | Mar 8 |
| Kappa | EPI_ISL_1716737 |  | Apr 6 |
| B.1.1.318 | EPI_ISL_2370918 |  | Feb 5 |

\* Eight, presumably Delta precursor isolates, were detected as follows: EPI\_ISL\_1716736 and EPI\_ISL\_2364992, carrying 19R, 156G, Δ157-158, 452R, 478K, 614G, 681R and 950N, were collected on March 23, 2021 and April 28, 2021 respectively; four isolates, harboring 19R, 156G, Δ157-158, 452R, 478K, 614G, and 681N (EPI\_ISL\_1921288, EPI\_ISL\_1921764, EPI\_ISL\_1921765, EPI\_ISL\_1921768), and two isolates with 19R, 158R, 452K, and 681R (EPI\_ISL\_1921165, EPI\_ISL\_1921739), were collected on April 26, 2021.

**SI Table 3.**
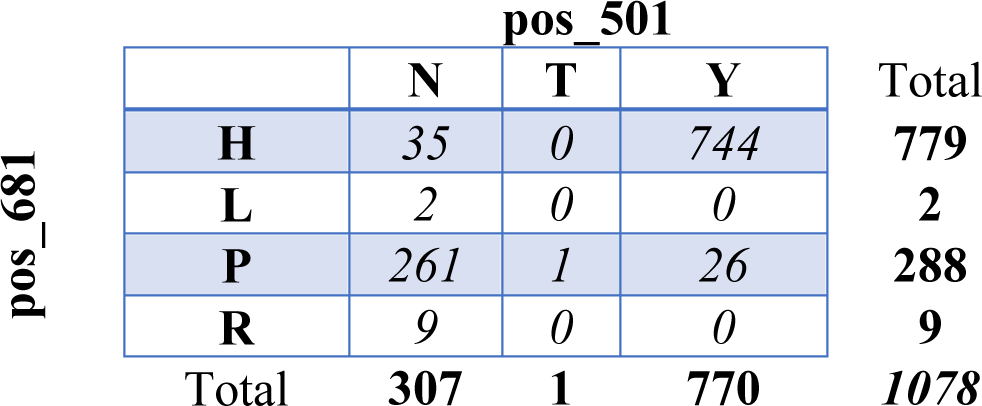
Exemplary 4×3 contingency table, one of the 14,365 r × c tables examined, showing the marginal (total row or total column), conditional (rows or columns) and joint (cells) distributions of the amino acid frequencies between functional positions 681 and 501 in the Greek dataset of SARS-CoV-2 spike proteins. The probability value in all three tests of independence for this table was ≤0.001, implying an association between specific combinations of residues at the two positions.

